# A host E3 ubiquitin ligase regulates *Salmonella* virulence by targeting an SPI-2 effector involved in SIF biogenesis

**DOI:** 10.1101/2022.08.05.502941

**Authors:** Kun Meng, Jin Yang, Juan Xue, Jun Lv, Ping Zhu, Liuliu Shi, Shan Li

## Abstract

*Salmonella* Typhimurium creates an intracellular niche for its replication by utilizing a large cohort of effectors, including several that function to interfere with host ubiquitin signaling. Although the mechanism of action of many such effectors has been elucidated, how the interplay between the host ubiquitin network and bacterial virulence factors dictates the outcome of infection largely remains undefined. Here we found that the SPI-2 effector SseK3 inhibits SNARE pairing to promote the formation of *Salmonella*-induced filament by Arg-GlcNAcylation of SNARE proteins, including SNAP25, VAMP8, and Syntaxin. Further study reveals that host cells counteract the activity of SseK3 by inducing the expression of the ubiquitin E3 ligase TRIM32, which catalyzes K48-linked ubiquitination on SseK3 and targets its membrane-associated portion for degradation. Hence, TRIM32 antagonizes SNAP25 Arg-GlcNAcylation induced by SseK3 to restrict SIF biogenesis and *Salmonella* replication. Our study reveals a mechanism by which host cells inhibit bacterial replication by eliminating specific virulence factor.

## Introduction

*Salmonella enterica serovar* Typhimurium (*S*. Typhimurium) is a facultative intracellular pathogen capable of infecting multiple hosts to cause salmonellosis (Acheson and Hohmann, 2001). *S*. Typhimurium encodes two type III secretion systems (T3SSs), SPI-1 and SPI-2 that inject a large cohort of effector proteins into host cells(Dos Santos et al., 2020). Whereas the function of SPI-1 primarily is to promote invasion of non-phagocytic intestinal epithelial cells to establish a nascent phagosome, effectors translocated by SPI-2 function to remodel the phagosome into a niche permissive for bacteria replication called the *Salmonella*-containing vacuole (SCV)(Lou et al., 2019). A number of SPI-2 effectors are dedicated to subverting the endolysosomal system, including the formation of a complex and highly dynamic structure termed *Salmonella*-induced filaments (SIFs)(Jennings et al., 2017; Knuff and Finlay, 2017). Although many pathogens have the ability to create bacteria-containing vacuoles (BCVs) to support their intracellular replication, the formation of the SIF network is unique to *S*. Typhimurium (Santos and Enninga, 2016). The ability of this pathogen to form SIF strongly correlates with its ability to cause diseases in animal infection models (Stein et al., 1996). It had been hypothesized that SIF structures facilitate the bacterium to gain access to nutrients and/or to evade host immunity (Liss et al., 2017; Noster et al., 2019). Thus, understanding the mechanisms of SIF biogenesis and maintenance will not only lead to a better appreciation of the *S*. Typhimurium pathogenesis and host cell biology but also provide clues for its disruption as novel interference strategies against salmonellosis.

Ubiquitin signaling plays an essential role in immunity by regulating various cellular processes, particularly protein turnover and vesicle trafficking(Dikic, 2017). Consistent with the existence of multiple mechanisms designed to sense and eliminate invading *S*. Typhimurium, the bacterium has evolved several virulence factors to counteract such surveillance by directly interfering with ubiquitin signaling. At least four Salmonella effectors have been shown to function as E3 ubiquitin ligases to modulate host immunity(Herhaus and Dikic, 2018). Among them, SopA is a HECT-like E3 ligase (Zhang et al., 2006) that modifies and targets TRIM56 and TRIM65, two host E3 ligases for degradation(Fiskin et al., 2017; Kamanova et al., 2016). The two IpaH-type E3 enzymes SspH1 and SspH2 target protein kinase 1 and NO1, respectively (Bhavsar et al., 2013; Haraga and Miller, 2006; Keszei et al., 2014). SlrP promotes host cell death by targeting thioredoxin and the Hsp40/DnaJ chaperone family associated with the endoplasmic reticulum (Bernal-Bayard et al., 2010; Bernal-Bayard and Ramos-Morales, 2009). In addition, the two deubiquitinases, AvrA and SseL have been found to regulate host immunity, particularly the NF-κB pathway(Hermanns and Hofmann, 2019; Mesquita et al., 2013; Ye et al., 2007). The fate of *S*. Typhimurium that escape the phagosome to reach the cytosol differs greatly from those residing in the membrane-bound vacuoles, these bacteria are first ubiquitinated by the E3 ligase RNF213, which strikingly directly recognizes and modifies the lipid A moiety of bacterial lipopolysaccharide (LPS)(Otten et al., 2021). Several other E3 ligases coordinate to build the ubiquitin coat on the bacterial surface to initiate xenophagy(Herhaus and Dikic, 2018), a process that was recently found to be inhibited by the effector SopF that ADP-ribosylates the ATP6V0C subunit of the v-ATPase complex (Xu et al., 2019).

Another cohort of *S*. Typhimurium effectors contribute to the development of the SCV by modulating vesicle trafficking(Galán, 2021). Although the cellular function of these effectors has been extensively studied (Tuli and Sharma, 2019), how the host cell counteracts their activity largely remains elusive. In the present study, we identified the host E3 ligase TRIM32 as a regulator for the activity of the SPI-2 effector SseK3. We found that SseK3 catalyzes Arg-GlcNAcylation on SNAP25 and restricts SIF biogenesis mediated by the SseK3-SNARE axis and that infection by *S*. Typhimurium induces the expression of TRIM32, which antagonizes the activity of SseK3 by ubiquitination-mediated degradation.

## Results

### SseK3 promotes SIF formation during *S.* Typhimurium infection

Effectors translocated by the SPI-2 are required for the formation of SIFs (Knuff and Finlay, 2017). Among these, the three SseK proteins, SseK1, SseK2 and SseK3, are arginine GlcNAc transferases that are important for *S.* Typhimurium virulence(Meng et al., 2020). To test whether any of these effectors is involved in SIF biogenesis or maintenance, we examined the phenotypes by infecting a cell line derived from HeLa that expresses GFP-VAMP8 with several relevant *S.* Typhimurium strains, including the wild-type, Δ*sseK1/2/3*, and Δ*ssaV* which is defective in the SPI-2. Samples were assessed for the integrity of SCV membrane and SIF formation by confocal microscopy. At 2 h post-invasion, the frequency of VAMP8-coated SCVs for strain Δ*sseK1/2/3* was similar to that of the wild-type and the Δ*ssaV* mutant, with rates at 81±2%, 83±6%, and 83.1±4%, respectively **(Figure S1)**. Thus, SseKs do not contribute to the formation of nascent SCVs, which is consistent with the observation that Arg-GlcNAcylation induced by members of SseK family does not occur until after 6 h post-infection(Meng et al., 2020). At 10 h post-invasion, we observed a significant decrease in the frequency of SIFs in cells infected with the Δ*sseK1/2/3* strain when compared to those infected with wild-type *S*. Typhimurium. Furthermore, expression of SseK3, but not its enzymatically inactive mutant SseK3_D226A/D228A_ restored the development of SIF to wild-type levels (**Figures 1A** **and 1B)**. These results indicate that SseK3 plays a role in promoting SIF formation during *S.* Typhimurium infection.

**Figure 1.**
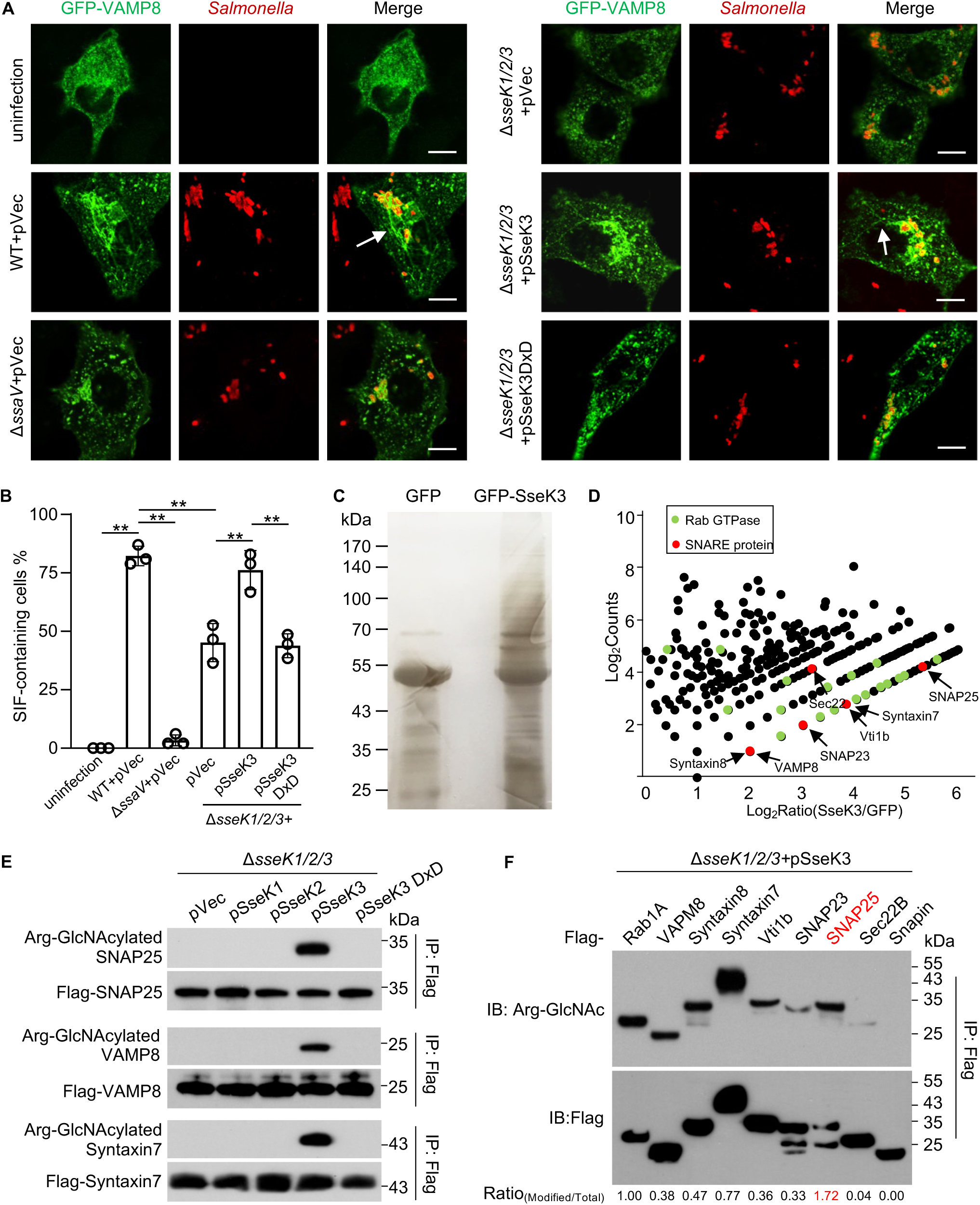
SseK3 promotes SIF formation by modifying SNARE proteins during *S.* Typhimurium infection. (A-B) SseK3 promotes SIF formation during *S.* Typhimurium infection. HeLa cells stably expressing EGFP-VAMP8 were infected with the indicated *S.* Typhimurium strains for 10 h and analyzed for SIFs. (A) Representative images showing the distribution of VAMP8 (green) and *S.* Typhimurium (red). Scale bar, 10 μm. (B) The rates of VAMP8-positive tubules for each sample are indicated. At least 50 cells were counted for samples from experiments done in triplicate. \**P*<0.05 (C) Detection of enriched Arg-GlcNAcylated proteins by silver staining. Lysates of 293T cells transfected to express GFP or GFP-SseK3 were subjected to immunoprecipitation with Arg-GlcNAc-specific antibodies, precipitates separated by SDS-PAGE were detected by silver staining. (D) Scatter plots of protein ratios as a function of their relative abundance. Proteins immunoprecipitated with an anti-Arg-GlcNAc antibody were subjected to LC-MS/MS analysis. The ratio was calculated as spectral counts in SseK3-transfected samples divided by those in GFP-transfected samples. Large ratios indicate preferential detection and modification in 293T cells transfected to express SseK3. Red dots correspond to SNARE proteins, and green dots correspond to Rab GTPase proteins. (E) SseK3 but not SseK1 or SseK2 modifies SNARE proteins during *S.* Typhimurium infection. 293T cells expressing the indicated Flag-tagged SNARE proteins were infected with relevant *S.* Typhimurium strains. Lysates immunoprecipitated with the Flag-specific antibody were detected by immunoblotting with the indicated antibodies. Similar results were obtained from at least three independent experiments. (F) SNARE proteins are modified by SseK3 during *S.* Typhimurium infection. 293T cells expressing the indicated Flag-tagged SNARE-associated proteins were infected with *S*. Typhimurium strain Δ*sseK1/2/3*(pSseK3). The level of Arg-GlcNAcylation was obtained by measuring band intensity of Arg-GlcNAcylated proteins to total protein using the image J software.

### SseK3 attacks SNARE proteins by Arg-GlcNAcylation

We next investigated the mechanism underlying SseK3-induced SIF development. Our previous studies have shown that SseK3 catalyzes Arg-GlcNAcylation on death domain-containing receptor proteins and members of the Rab small GTPases(Meng et al., 2020; Pan et al., 2020). Yet, despite extensive efforts, we did not detect SseK3-induced modification of most of the Rabs involved in SIF formation, including Rab11, Rab9, and Rab7 in cells infected with *S.* Typhimurium (Meng et al., 2020). We thus attempted to identify SseK3 targets involving in SIF biogenesis by Arg-GlcNAc (Pan et al., 2014) followed by mass spectrometry analysis (**Figure 1C**). KEGG analysis of the putative SseK3 modified proteins revealed five significantly enriched pathways. Among them, nineteen proteins are involved in SNARE interactions in the vesicular transport pathway, including SNAP23, SNAP25, VAMP8, Vti1b, Syntaxin7, Syntaxin8, and Sec22b (**Figure 1D**, red dots). Further comparative analyses between SseK3 samples and controls led to the identification of several GTPase proteins that had been previously identified (green dots, e.g., Rab1 and Rab8), demonstrating the effectiveness of this strategy (**Figure 1D**). Complementation experiments with SseK1, SseK2 or SseK3 showed that SseK3 was the sole enzyme responsible for modifying SNARE proteins during infection (**Figure 1E**). Further experiments established that several SNARE proteins, including VAMP8, Syntaxin7, Syntaxin8, Vti1b, Sec22b, SNAP23, and SNAP25 are modified by SseK3 in cells infected with *S.* Typhimurium (**Figure 1F**). Among these, SNAP25 was most robustly modified by SseK3, suggesting that this protein is its preferred substrate (**Figures 1F** **and S2)**. Taken together, these results indicate that SNARE proteins are new cellular targets of SseK3.

### SseK3 induces GlcNAcylation of SNAP25 on Arg30 and Arg31, two residues important for its interaction with VAMPs

SNARE proteins share a common coiled-coil motif of 60–70 residues essential for membrane fusion (**Figure 2A**)(Harbury, 1998). By deletion mutagenesis, we found that the amino-terminal domain (1-90 aa, NTD) containing the coiled-coil motif of SNAP25 can be modified by SseK3 in cells infected with *S.* Typhimurium (**Figure 2B**). To precisely map the modification site(s), we affinity-purified Flag-SNAP25 from 293T cells co-transfected with either wild-type SseK3 or empty vector. By tandem MS (MS/MS) analysis, we detected one GluC-LysC peptide ^28^STRRMLQLVEE^38^ with a mass shift of 406 Da only in samples from cells that co-expressed SseK3. The 406 Da increase in mass corresponds to the attachment of two GlcNAc moieties. Quantitation analysis revealed that approximately 75% of the peptides were modified **(Figure S3)**. MS/MS analyses assigned the modification sites to Arg30 and Arg31 (**Figure 2C**), which were further verified by mutagenesis analysis. Modification signals were no longer detected in samples expressing the SNAP25 R30K/R31K mutant (**Figure 2D**). Importantly, both Arg30 and Arg31 are located within the coiled-coil domain of the *t*-SNARE, which is involved in SNAP25 self-association and in interactions with its binding proteins (**Figure 2E**). To determine the impact of SseK3 on the binding of SNAP25 to its interacting partners, we used mass spectrometry to analyze proteins pulled down by SNAP25 under conditions with and without SseK3, which revealed that VAMP8 was not present in the pulldown products from samples that expressed SseK3 (**Figure 2F**). This phenomenon can be recapitulated by experiments in which such binding was detected by immunoblotting of samples that expressed enzymatically inactive SseK3 (**Figure 2G**).

**Figure 2.**
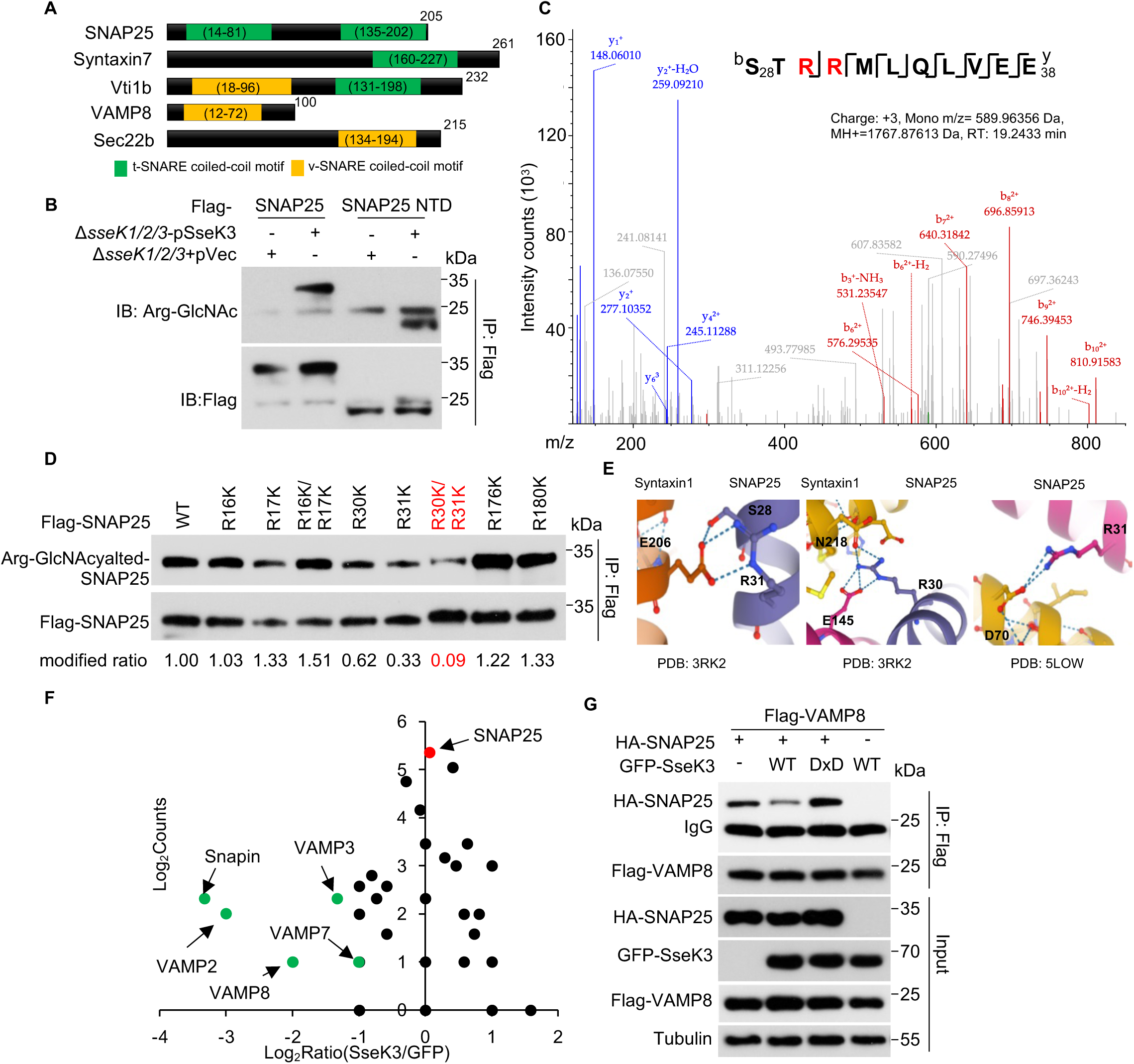
SseK3 interferes with the interactions between SNAP25 and VAMPs. (A) Schematic representation of the domain structure of known SNARE proteins. Each protein was represented by a rectangular bar and the localization of the t-SNARE coiled-coil domains and the v-SNARE coiled-coil domains are shown. (B) SseK3 modifies SNAP25 within the N-terminal domain during *S.* Typhimurium infection. Cells expressing the indicated domains of SNAP25 were infected with the indicated *S.* Typhimurium strains and the modification was detected by immunoblotting. (C) Determination of modification sites by mass spectrometric analysis. Shown is the MS/MS spectrum of modified peptide ^28^STRRMLQLVEE^38^. The fragment ions b_6_ to b_10_ have a mass increase of 406 Da corresponding to the addition of two GlcNAc while y_1_ to y_6_ fragments lack such a mass shift, indicating that GlcNAcyaltion occurs on Arg30 and Arg31. (D) Validation of Arg30 and Arg31 as the main modification sites of SNAP25 by SseK3. 293T cells expressing the indicated Flag-tagged SNAP25 mutants were infected with *S*. Typhimurium strain Δ*sseK1/2/3*(pSseK3), samples lysed were immunoprecipitated with anti-Flag beads and detected with antibodies specific for the Arg-GlcNAcylation. The levels of modification were quantitated by measuring the ratio of band intensity for modified and total proteins with Image J. (E) 3D structure visualization of Arg30 and Arg31 in SNAP25 (PBD number: 5LOW) and in the SNAP25-syntaxin1 complex (PBD number:3RK2). Note the role of the two residues in interactions between these two proteins. (F) Quantification of SNAP25-binding proteins in cells expressing SseK3. Proteins that potentially bind SNAP25 were obtained by IP from cells transfected to express SseK3 were analyzed by mass spectrometry, cells transfected with the vector were used as controls. Scatter plots of protein ratios as a function of their relative abundance (denoted by MS/MS spectral counts). The ratio is calculated as spectral counts in SseK3 transfected samples divided by those in controls. Lower ratios indicate decreased binding efficiency with SNAP25. Green dots correspond to Snapin and VAMPs, and the red dot corresponds to immunoprecipitated SNAP25. Results shown are a representative of three independent experiments with similar results. (G) SseK3 interferes with the interactions between SNAP25 with VAMP8. Lysates of cells transfected to express the indicated protein combinations were subjected to IP with Flag-specific antibody. The products were detected for the presence of the binding partners by immunoblotting. Similar results were obtained in three independent experiments.

### SseK3 limits the size of SNAP25-decorated infection-associated macropinosomes (IAMs) during late stages of infection

SNAP25 is enriched on the fluid-filled infection-associated macropinosome (IAM) (Stévenin et al., 2021), and it plays a crucial role in homotypic fusion among IAMs and their heterotypic fusion with SCVs, an event that is required for the expansion of the bacterial phagosome shortly after phagocytosis (Stévenin et al., 2019). To explore the possibility that SseK3-induced GlcNAcylation may interfere with its ability to promote such fusion events and the expansion of the SCV, we measured the dimension of the SNAP25-containing vacuoles (SNAP25-CVs) that represent SCVs (with bacteria) and IAMs (without bacteria), respectively. Our results indicate that at 0.5 h and 2 h infection time, there was no significant difference between the diameter of SNAP25-CVs found in cells infected with wild-type bacteria and the Δ*sseK1/2/3* mutant. Intriguingly, when the infection has proceeded for 6 h, the size of the SNAP25-CVs was significantly smaller in cells infected with the wild-type strain than that found in cells infected with the Δ*sseK1/2/3* mutant (**Figure 3A**). Such difference disappeared when SseK3 was expressed in strain Δ*sseK1/2/3* from a plasmid (**Figures 3B** **and 3C)**. In agreement with these results, in cells transfected to express SseK3 prior to infection, the size of vacuoles formed by strain Δ*sseK1/2/3* became smaller (**Figures 3D** **and 3E)**. Furthermore, signals of protein Arg-GlcNAcylation induced by SseK3 in infected cells displayed clear co-localization with SNAP25 on bacterial vacuoles (**Figure 3D**). Thus, SseK3 limits the size of the SNAP25-containing IAM vacuole when the infection has proceeded for 6 h. Given the fact that SseK3 attenuates the interaction between SNAP25 and VAMPs (**Figure 3F**), such alternations in the size of the vacuoles may be caused by the inhibition of fusion among IAMs and between IAMs and SCVs. Because SNAREs are essential for SIF formation (Kehl et al., 2020), it is likely that SseK3 interferes with SNARE paring to impact SCV size and SIF formation at the late stage of infection (**Figure 1A**).

**Figure 3.**
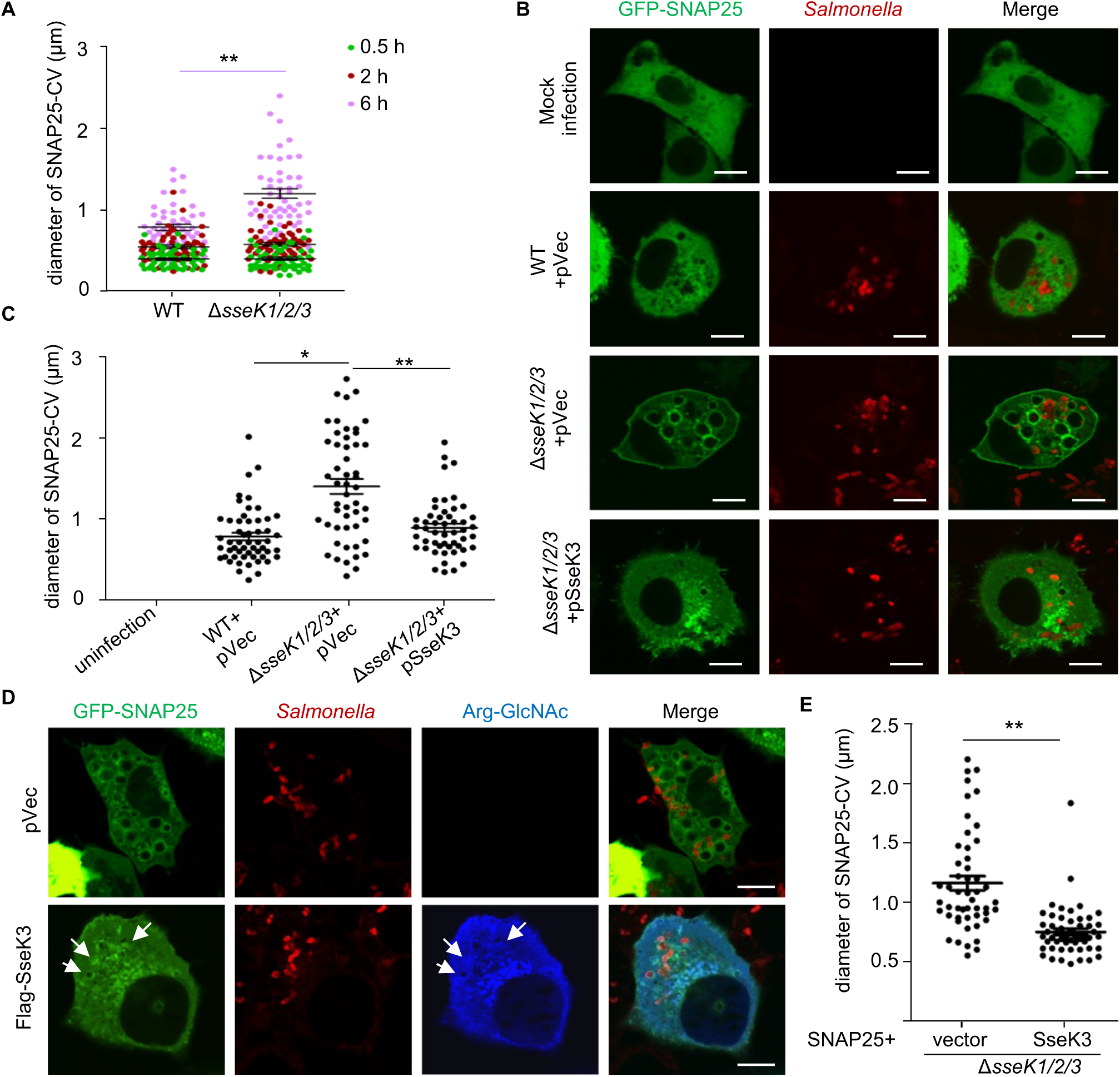
SseK3 limits the size of SNAP25-labeled SCVs. (A) The effects of the SseK family proteins on the size of SNAP25-containing vacuole (SNAP25-CV) during *S.* Typhimurium infection. HeLa cells transfected to express GFP-SNAP25 were infected with the indicated *S.* Typhimurium strains. The diameter of SNAP25-CV was measured at the indicated time points. (B-C) The effects of SseK3 on the size of SNAP25-CV. HeLa cells transfected to express GFP-SNAP25 were infected with the indicated *S.* Typhimurium strains for 6 h. The distribution of SNAP25 (green) and *S.* Typhimurium (red) are shown in B. Statistics of the diameter of SCVs positive for GFP-SNAP25 are shown in C. (D-E) Ectopic expression of SseK3 limits the size of SNAP25-CV. HeLa cells transfected to express GFP-SNAP25 and Flag-SseK3 were infected with the indicated *S.* Typhimurium strains for 6 h. The distribution of GFP-SNAP25 (green), *S.* Typhimurium (red), and Arg-GlcNAcylated proteins (blue) were shown (D). Statistics of the diameter of the SNAP25 positive SCVs are shown in E. Arrow-heads indicate the co-localization of SNAP25 and Arg-GlcNAcylation on the vacuoles. At least 30 cells in A, C and E were analyzed for each experiment. Scale bar, 10 μm. \**P*<0.05, \*\**P*<0.01.

### Expression level of the host ubiquitin E3 ligase TRIM32 is induced upon *S*. Typhimurium infection

To determine the mechanism the host may employ to counteract the activity *S*. Typhimurium effectors, we re-analyzed the available transcriptomic data of host cells infected with strain SL1344 (Avraham et al., 2015). These efforts led to the identification of *trim32* that codes for an E3 ubiquitin ligase as one of the significantly induced genes in response to *S*. Typhimurium challenge (**Figure 4A**). Further analyses by quantitative real-time PCR (qRT-PCR) and immunoblotting confirmed that *trim32* is induced at both mRNA and protein levels upon bacterial challenge (**Figures 4B** **and 4C**). Moreover, overexpression of TRIM32 led to a significant decrease in the frequency of the intracellular SIF structures found in cells infected with *S*. Typhimurium (**Figures 4D** **and 4E**).

**Figure 4.**
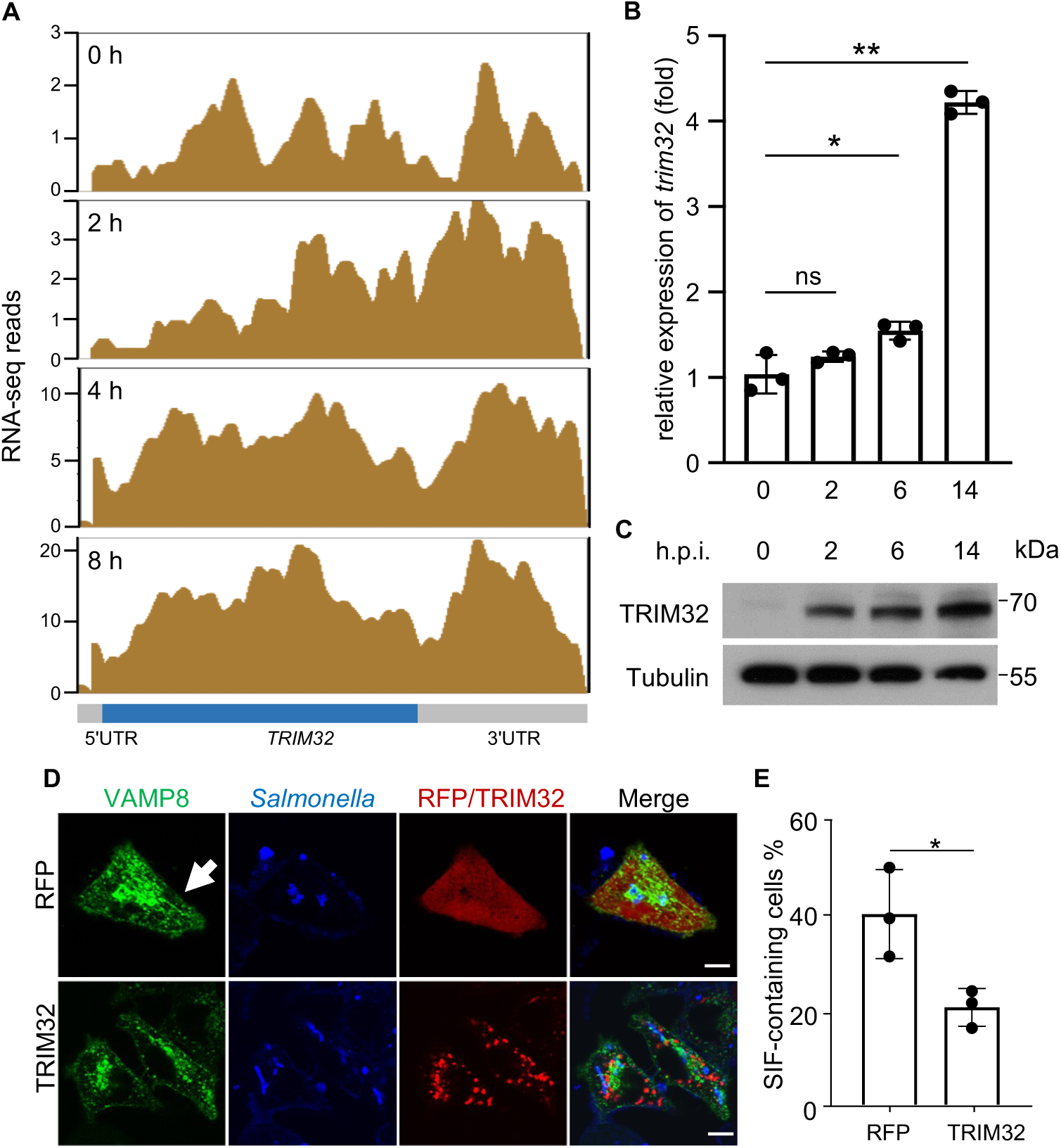
Expression of the host E3 ligase gene *trim32* is induced in response to *S.* Typhimurium infection. (A) Diagrams of the *TRIM32* locus based on RNA sequencing reads. Enriched RNA-seq signals (Avraham et al., 2015) visualized by Integrated Genome Browser are representative of three independent experiments. UTR, the untranslated region. (B) qRT-PCR detection of *trim32* expression during *S.* Typhimurium infection. HeLa cells infected with *S.* Typhimurium strain SL1344 for the indicated time points were probed for the mRNA levels of *trim32*. The statistical data are expressed as means± SD from three independent experiments. \**P*<0.05, \*\**P*<0.01. (C) The induction of TRIM32 in response to *S*. Typhimurium infection measured by detecting protein. HeLa cells infected with *S*. Typhimurium strain SL1344 for the indicated time points were probed with TRIM32-specific antibodies. Tubulin was detected as a loading control. Data shown are one representative from three independent experiments with similar results. (D-E) The effects of TRIM32 on SIF biogenesis. HeLa cells transfected to express GFP-VAMP8 and the indicated proteins for 12 h were infected with the indicated *S.* Typhimurium for 10 h. (D) The distribution of GFP-VAMP8 (green), *S.* Typhimurium (blue), and RFP or RFP-TRIM32 was shown. (E) Quantitation of cells showing VAMP8-positive tubules is indicated. At least 50 cells were counted for each experiment and the statistical data shown are from three independent experiments. Arrow-heads indicate the SIF structure. Scale bar, 10 μm. \**P*<0.05

### TRIM32 interacts with and ubiquitinates SseK3

An earlier study has demonstrated interactions between TRIM32 and SseK3 (Yang et al., 2015). Yet, SseK3 does not detectably modify TRIM32 by Arg-GlcNAcylation, and the physiological significance of this interaction is unknown. TRIM32 consists of an amino-terminal RING domain, a type II B-box domain, a coiled-coil domain, and a carboxyl NHL domain. To determine which of these regions is important for its interaction with SseK3, we generated a series of TRIM32 deletion mutants and examined their ability to bind the effector. Results from these experiments indicate that the NHL domain is essential for SseK3 binding (**Figures 5A** **and 5B)**.

**Figure 5.**
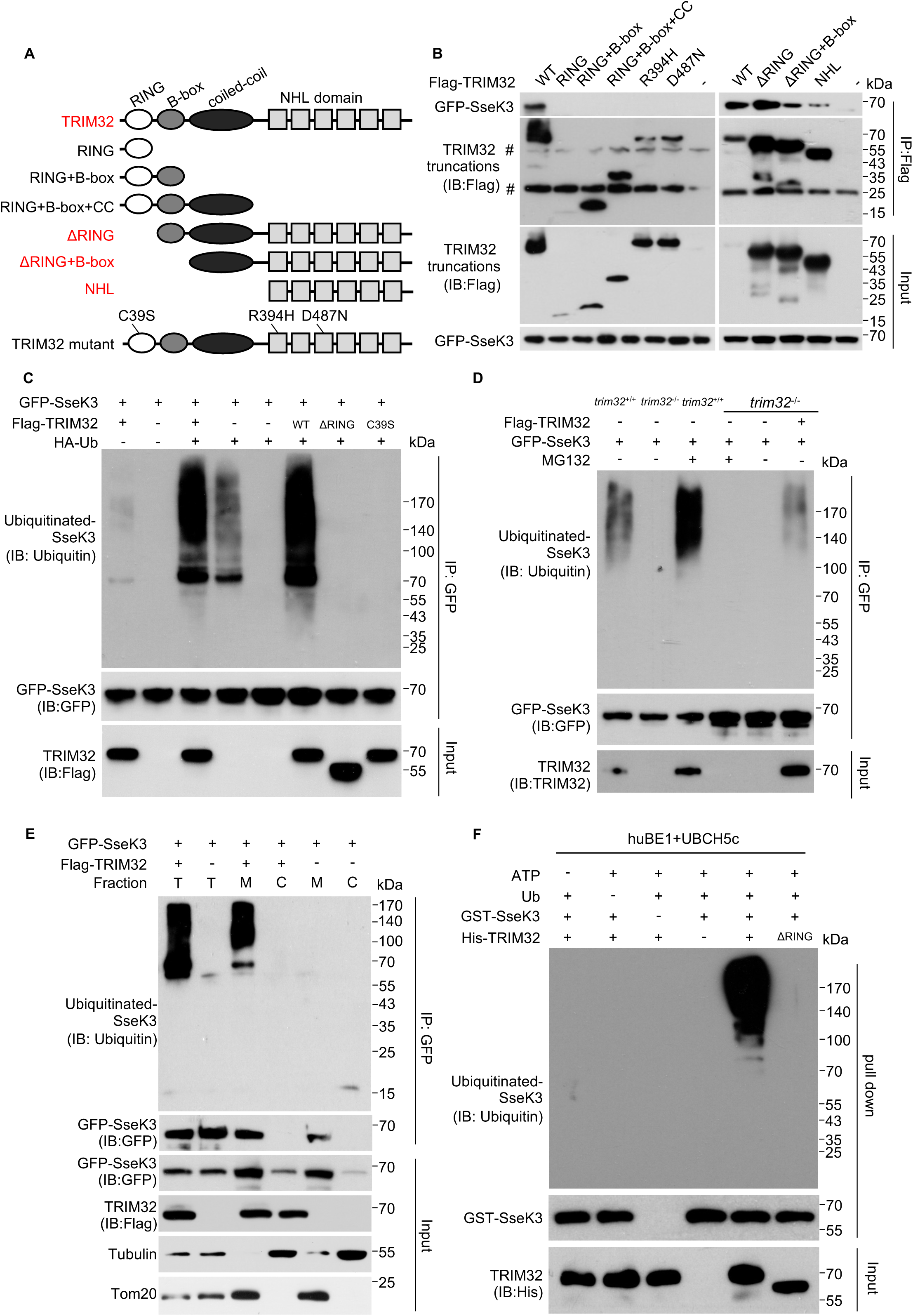
TRIM32 interacts with and ubiquitinates SseK3. (A) A schematic representation of TRIM32 domain structure and the several TRIM32 mutants used in this study. TRIM32 truncation mutants that retain the ability to interact with SseK3 are shown in red. (B) Mapping the domain important for TRIM32 to interact with SseK3. Flag-tagged full-length or several deletion mutants of TRIM32 was coexpressed GFP-SseK3 in 293T cells and the interactions were determined by immunoprecipitation with beads coated with the Flag-specific antibody. Binding was detected by immunoblotting with GFP-specific antibodies. # marks the IgG in the IP blot. (C) Overexpression of wild-type but not the mutant TRIM32 promotes ubiquitination of SseK3. 293T cells transfected to express GFP-SseK3 and TRIM32 or its mutants in the presence or absence of HA-ubiquitin. 18 h after transfection, co-immunoprecipitation was performed with anti-GFP antibodies, followed by standard immunoblotting analysis with the indicated antibodies. (D) TRIM32 is required for ubiquitination of SseK3. The indicated cell lines, *trim32*^+/+^, *trim32*^-/-^ and *trim32*^-/-^ complemented with TRIM32 were transfected to express GFP-SseK3 for 16 h. Cells were treated with or without 25 μM MG132 for 12 h prior to probing for the ubiquitination levels of GFP-SseK3 by immunoblotting. (E) TRIM32-mediated ubiquitination of SseK3 occurs at the membrane components of SseK3. 293T cells were transfected with the indicated plasmids. Total membrane and cytosol proteins were isolated and immunoblotted with the corresponding antibodies. T: total protein. M: membrane fraction. C: cytoplasmic fraction. (F) TRIM32 catalyzes SseK ubiquitinatination *in vitro*. Recombinant His-TRIM32, GST-SseK3, ubiquitin, E1 (huBE1), and E2 (UBCH5c) were added as indicated for ubiquitination assays. After GST pull-down, ubiquitin-conjugated proteins were detected by immunoblot with a ubiquitin-specific antibody. The input levels of TRIM32 proteins were detected by immunoblots. Data shown are a representative of three independent experiments with similar results.

TRIM32 is an E3 ubiquitin ligase that contains a RING finger domain. Therefore, we examined whether TRIM32 can catalyze ubiquitination on SseK3. Co-expression of TRIM32 with SseK3 and HA-ubiquitin led to ubiquitination of the effector. In contrast, no ubiquitination signal was detected in samples from cells transfected to express TRIM32ΔRING lacking the RING domain or the enzymatically inactive mutant TRIM32(C39S) in which the active cysteine was mutated to alanine (Kehl et al., 2020) (**Figure 5C**). In agreement with these results, knocking-out *trim32* diminished SseK3 ubiquitination in a way that can be complemented with the wild-type gene but not the C39S mutant (**Figure 5D**). Furthermore, treatment of the cells with the proteasome inhibitor MG132 considerably enhanced SseK3 ubiquitination (**Figure 5D**). SseK3 is a Golgi-located protein(Meng et al., 2020), and we found that TRIM32-mediated ubiquitination occurred at the membrane components of SseK3 (**Figure 5E**). Finally, in biochemical assays, inclusion of TRIM32 in reactions containing huBE1 (E1), UBCH5c (E2), Ub, and ATP led to robust SseK3 ubiquitination. Such modification did not occur in reactions receiving TRIM32ΔRING (**Figure 5F**).Thus, TRIM32 ubiquitinates SseK3 by its NHL domain recognition.

### TRIM32 catalyzes K48-linked ubiquitination on SseK3 and targets its membrane-associated portion for degradation

There are seven lysine residues (K6, K11, K27, K29, K33, K48, and K63) in the ubiquitin molecule, each dictates the formation of a specific type of polyubiquitin chain that determines the biological consequence of the modification (Walczak et al., 2012). To further understand the implications of TRIM32-induced ubiquitination on the activity of Ssek3, we evaluated the chain type of the ubiquitin polymers on SseK3 using a series of lysine mutants of ubiquitin. Robust ubiquitination of SseK3 was observed in reactions receiving wild-type or K48-only ubiquitin (a mutant in which all Lys residues except for Lys48 have been replaced with Arg). Conversely, in reactions containing the K48R ubiquitin mutant, no ubiquitination was detected (**Figure 6A**). Polyubiquitination by K48 chain type typically results in protein degradation by the ubiquitin-proteasome system (UPS)(Clague and Urbé, 2010), we then investigated the stability of SseK3 *in cellulo*. Chase experiments with cycloheximide (CHX) showed that the levels of SseK3 associated with the membrane (M-SseK3) were markedly reduced in a manner that can be blocked by MG132 (**Figure 6B**). These results strongly suggest that SseK3 was degraded via the UPS after being ubiquitinated by TRIM32.

**Figure 6.**
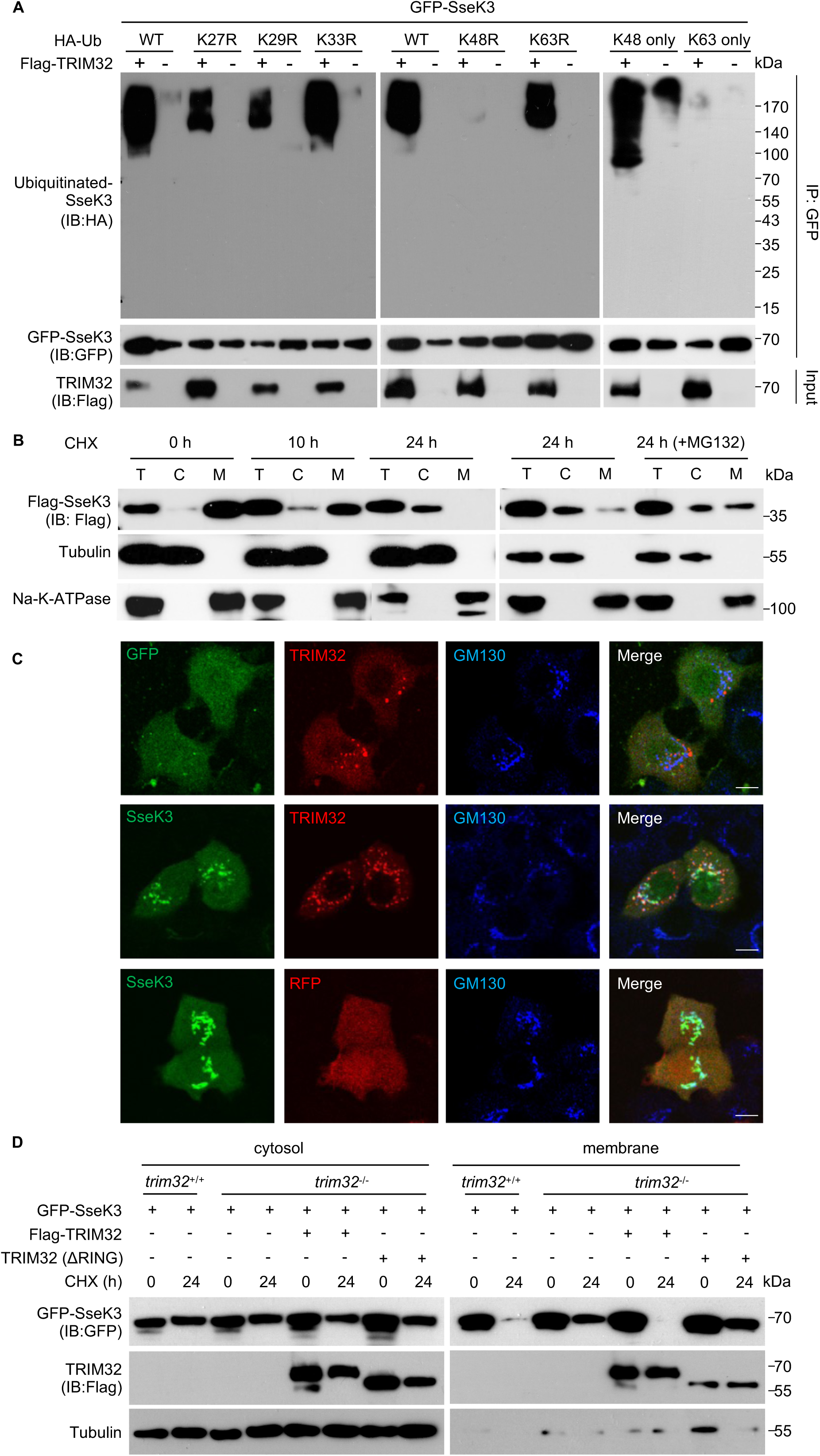
TRIM32 catalyzes K48-linked ubiquitination to degrade membrane-associated SseK3. (A) TRIM32 catalyzes K48-linked polyubiquitination of SseK3. 293T cells transfected to express GFP-SseK3 and the indicated proteins. Lysates of the samples were subjected to immunoprecipitation and immunoblot analysis with the indicated antibodies. (B) The stability of SseK3 probed by the CHX pulse chase-assay. 293T cells transfected to express Flag-SseK3 were treated with CHX for the indicated time points. Cells were treated with 25 μM MG132 for 12 h before cell lysis (B). Total membrane and cytosol proteins were isolated and immunoblotted with the corresponding antibodies. M: membrane fraction. C: cytoplasmic fraction. (C) Co-localization of TRIM32 and SseK3 on the Golgi apparatus in HeLa cells. HeLa cells transfected to express the indicated proteins were fixed with 4% paraformaldehyde and analyzed by confocal microscopy. The distribution of SseK3 or GFP (green), RFP-TRIM32 or RFP (red), and GM130 (blue) was shown. Scale bar, 20 µm. (D) The effects of TRIM32 on the stability of SseK3. *trim32*^+/+^, *trim32*^-/-^, and *trim32*^-/-^ cells complemented with TRIM32 each was transfected to express GFP-SseK3. Samples were treated with 100 μg/mL CHX for 24 h and were separated into membrane and cytosolic fractions. The presence of the relevant proteins in these fractions was probed by immunoblotting with the appropriate antibodies. Data shown are a representative of three independent experiments with similar results.

SseK3 has been shown to localize to the *cis*-Golgi apparatus(Meng et al., 2020). Importantly, RFP-TRIM32 also extensively targets to the Golgi apparatus when co-expressed with GFP-SseK3 in HeLa cells (**Figure 6C**). We therefore examined whether TRIM32 induces degradation of SseK3. Our results indicate TRIM32 indeed causes the reduction of SseK3 that is associated with membrane (**Figure 6D**). Consistently, knockout of *trim32* led to elevated SseK3 in the membrane fraction, which can be reversed by expressing TRIM32 but not the ΔRING mutant. In contrast, the protein level of cytosol fraction of SseK3 (C-SseK3) was largely unaffected by TRIM32 (**Figure 6D**). Thus, TRIM32 induces K48-linked ubiquitination on SseK3, which results in its degradation, particularly for protein that is associated with the membrane.

### TRIM32 antagonizes SNAP25 Arg-GlcNAcylation induced by SseK3 to restrict SIF biogenesis and *Salmonella* replication

The observation that TRIM32 ubiquitinates SseK3 and targets it for degradation suggests that this E3 ubiquitin ligase regulates the function of the effector. Indeed, we found that SseK3 injected into host cells by S. Typhimurium interacted with TRIM32 (Figure 7A). As expected, the level of GlcNAcylated SNAP25 decreased in samples expressing TRIM32 but not the mutant lacking the RING domain (Figure 7B). In line with these observations, overexpression of TRIM32 diminished the ability of SseK3 to promote SIF formation (**Figures 7C** **and 7D)**. Considering that both SseK3 protein and SIF structure are required for *Salmonella* intracellular survival within macrophages(Meng et al., 2020; Singh et al., 2018), we next determined the effects of TRIM32 on bacterial replication in macrophage cells. We synthesized three pairs of siRNA and found that the 2nd pair had the best down-regulation effect (**Figure 7E**). As expected, *trim32* knockdown efficiently facilitated bacterial replication of SseK3-expressing-*Salmonella*, but not S. Typhimurium lacking *sseKs* in RAW264.7 macrophage cells (**Figure 7F**). Together, these results indicate that TRIM32 functions to restrict the activity of SseK3 by targeting it for proteasome degradation, thus lowering Arg-GlcNAcylation on SNAP25, which ultimately causes a reduction in SIF formation and bacterial virulence (**Figure 8**).

**Figure 7.**
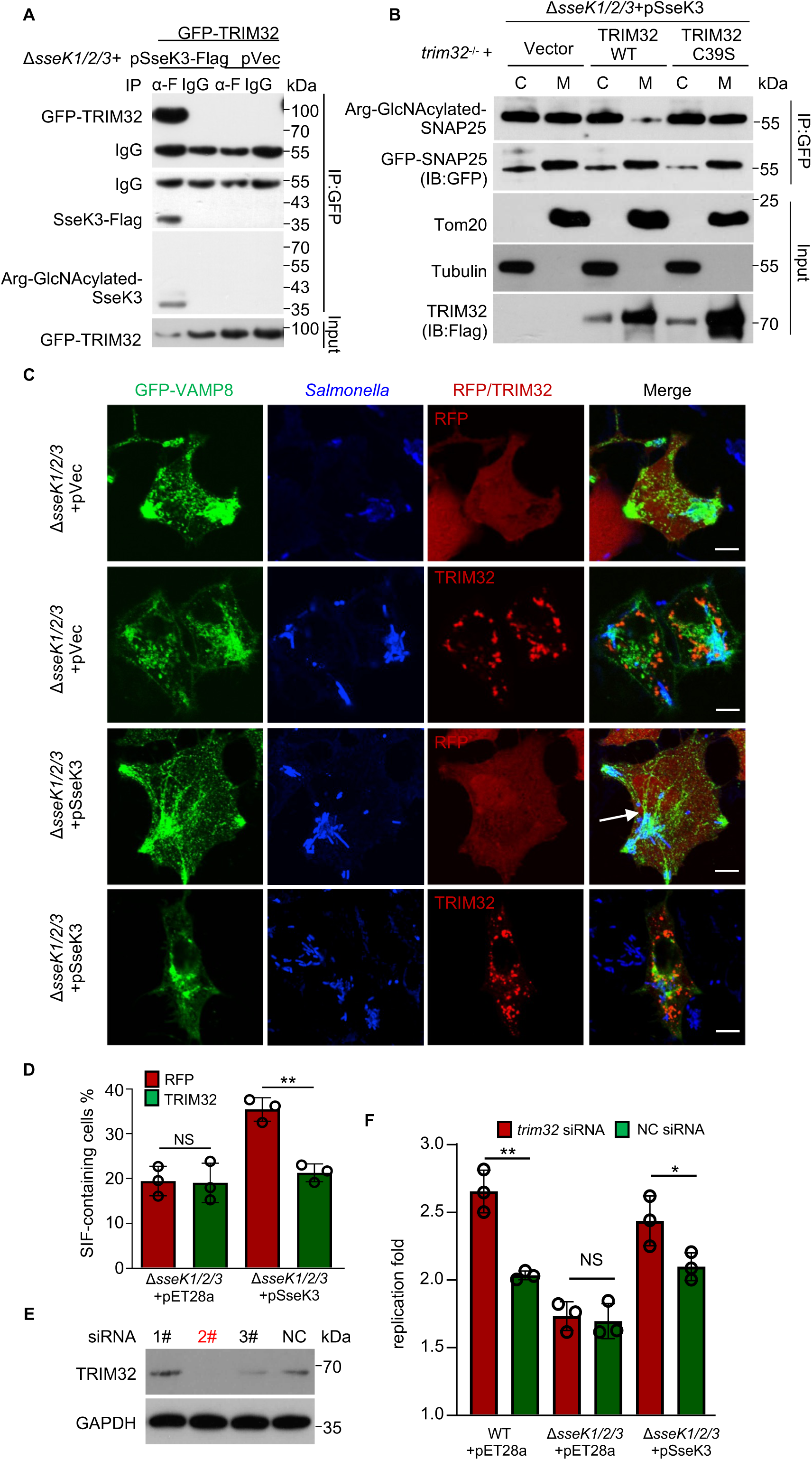
TRIM32 antagonizes SseK3-catalyzed-Arg-GlcNAcylation on SNAP25 and restricts SseK3-SNARE-mediated SIF biogenesis and *Salmonella* replication. (A) TRIM32 targets SseK3 during *S.* Typhimurium infection. 293T cells transfected to express GFP-TRIM32 were infected with the indicated bacterial strains for 16 h. The interactions between TRIM32 and SseK3 were detected by immunoprecipitation. (B) The effects of TRIM32 on the SseK3-catalyzed-Arg-GlcNAcylation on SNAP25 during *S.* Typhimurium infection. *trim32*^-/-^ cells transfected to express GFP-SNAP25 and the indicated proteins were infected with strain Δ*sseK1/2/3*(pSsek3) for 10 h. Chloramphenicol was added to inhibit the bacteria protein synthesis for another 12 h. Lysed cells were separated into soluble and membrane fractions and the presence of SseK3 was probed by immunoblotting. Data shown are a representative of three independent experiments with similar results. (C-D) The effects of TRIM32 on SseK3-SNARE-mediated SIF biogenesis. HeLa cells transfected to express GFP-VAMP8 and the indicated proteins for 12 h were infected with the indicated *S.* Typhimurium strains for 10 h. (C) The distribution of VAMP8 (green), *S.* Typhimurium (blue), and RFP or RFP-TRIM32 was determined by confocal microscopic analysis. Arrow-heads indicate the SIF structure. Scale bar, 10 μm. (D) The rates of cells showing VAMP8-positive tubules are indicated. At least 50 cells were counted for each sample done in triplicate and statistic data shown are from three independent experiments. \*\**P*<0.01. NS: not significant. (E-F) Effects of *trim32* knockdown on *Salmonella* replication in macrophage cells. (F) Knockdown efficiency of *trim32* siRNA was detected by immunoblotting. (F) RAW264.7 cells were transfected with 2# siRNA for *trim32* for 48 h, and then subjected to infection with the indicated *S*. Typhimurium at a multiplicity of infection of 10. Fold replication was determined by comparing bacterial counts at 2 and 24 h post-infection. Results shown are mean values ± SD (error bar) from three independent experiments. \**P*<0.05. \*\**P*<0.01. NS: not significant.

**Figure 8.**
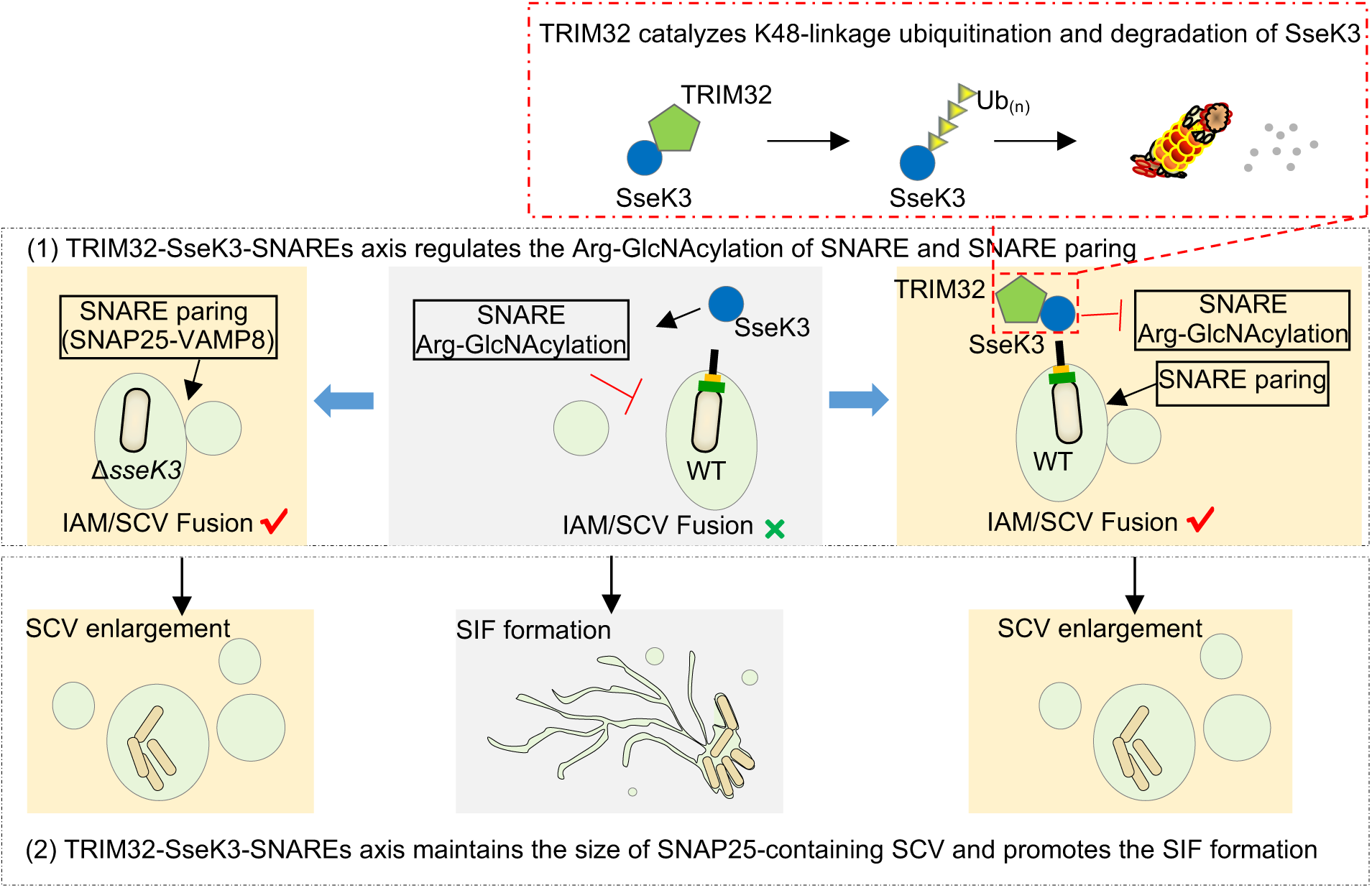
A schematic diagram of the regulation of *S*. Typhimurium intracellular lifecycle by SseK3 and TRIM32.

## Discussion

The outcome of bacterial infection is dictated by intimate interactions between host factors and virulence factors(Tripathi-Giesgen et al., 2021). Several members of the TRIM E3 ubiquitin ligase family have been found to be involved in regulating pathogen infection, including some that confer resistance to the virus by directly targeting viral proteins (Koepke et al., 2021; Lazzari and Meroni, 2016). TRIM56 and TRIM65 have been shown to be targeted by SopA, a HECT-like E3 ligase(Kamanova et al., 2016). Our discovery of TRIM32 as a host factor that functions to control *S.* Typhimurium virulence has expanded the role of these E3 ubiquitin ligases in defense against bacterial infection.

Importantly, an earlier study has shown that Trim32 plays a role in defending against *S.* Typhimurium infection because *trim32*^-/-^ mice are more sensitive to inflammatory death caused by this bacterium (Yang et al., 2017). In this study, several lines of evidence show that TRIM32 is involved in the defense against *S.* Typhimurium by attacking the SPI-2 effector SseK3. First, the expression of TRIM32 and SseK3 are synchronously upregulated. SseK3 belongs to an SPI-2 effector and begins to catalyze Arg-GlcNAcylation after 6 h post-infection(Meng et al., 2020). Both *trim32* mRNA and TRIM32 protein levels were significantly induced at this time (**Figure 4**). Second, TRIM32 interacts with SseK3 expressed by transfected or injected into host cells by *S.* Typhimurium via its carboxyl NHL repeats. Third, TRIM32 catalyzes K48-type polyubiquitin chains on SseK3 and targets its membrane-associated SseK3 for degradation. Fourth, overexpression of this E3 ligase led to less SIFs formation during *S.* Typhimurium infection, and *trim32* knockdown facilitated *Salmonella* replication within macrophage cells.

Our findings have also provided novel insights into the role of effector-induced Arg-GlcNAcylation in *S.* Typhimurium infection. Additional proteins potentially modified by members of the SseK have been identified in cells ectopically expressing the effectors or in cells infected with *S.* Typhimurium strains expressing specific effectors (Meng et al., 2020; Newson et al., 2019). These results are consistent with observations made in an earlier RNAi screen which revealed that the canonical mammalian late endo-/lysosomal vesicle fusion machinery (SNARE and Rab GTPase) is involved in SIF biogenesis(Kehl et al., 2020). Our finding that multiple coiled-coil containing-domain proteins are Arg-GlcNAcylated in infected cells suggests that SseKs may act coordinately to target endomembrane components to facilitate *S.* Typhimurium replication. In particular, SseK3 appears to directly participate in this process by imposing Arg-GlcNAcylation on SNAP25, leading to inhibition of its paring with VAMP8 and the restriction of the expansion of SCVs. This activity of SseK3 also causes a reduction in the size of macropinosomes, but the physiological consequence of such reduction is not clear. A recent screen using proximity labeling (BioID) found that SPI-2 effectors SseG, SopD2, PipB2, and SifA potentially interact with proteins in the SNARE complex (D’Costa et al., 2019). Interestingly, these effectors have been suggested to be involved in SIF formation(Knuff and Finlay, 2017). It is likely that multiple effectors, including SseK3 coordinate to regulate the dynamics of SNARE pairing to promote SIF biogenesis during *S.* Typhimurium infection. Future research aiming at dissecting the potential interplay among these effectors and how each is temporally and spatially regulated to ensure successful infection will yield more insights into their roles in the intracellular lifecycle of the pathogen.

A majority of proteins anchored in membrane-containing organelles have been reported to be degraded via the cytosolic proteasome system(Guo, 2022; Ramachandran et al., 2018). The spatial separation between substrate selection and degradation requires either membrane-anchored substrates extraction from the membrane or recruitment of 26S proteasomes to the membrane. A well-studied example is the endoplasmic reticulum (ER)-associated protein degradation (ERAD) pathway. Ubiquitinated proteins on ER are extracted from the membrane by Cdc48p/p97 complex and conveyed to the proteasome(Hwang and Qi, 2018). Besides, FKBP38, residing in the ER and mitochondrial membranes, functions to anchor the 26S proteasome to the organellar membrane(Nakagawa et al., 2007). Recently, proteasomes have been also reported to be constitutively associated with the Golgi membranes by PSMD6 and mediate the degradation of the Golgi protein GM130(Eisenberg-Lerner et al., 2020). SseK3 is a Golgi-located protein, and we here show that membrane counterparts of SseK3 are ubiquitinated and degraded in a proteasome-dependent manner. The mechanism is likely similar to GM130, and further investigations are needed.

Based on our results, we propose a model for the regulatory role of the TRIM32-SseK3-SNARE-SIF axis in *S.* Typhimurium infection. A few minutes after bacterial entry, the SCV increases in size through fusions with the macropinosomes. The *t*-SNARE protein SNAP25 localized on IAMs and v-SNARE VAMP8 are recruited to the SCV. The fusion between the SCV and IAMs allows the former to expand in size. As the infection has proceeded to late stages (>6h), SseK3 injected by the SPI-2 attacks SNAP25 by Arg-GlcNAcylation, leading to ablation of SNAP25-VAMP8 paring and the SNARE fusion events, which may contribute to SIF formation. Meanwhile, infected cells sense the presence of *S.* Typhimurium and induce the expression of *trim32* by a yet unrecognized mechanism. Elevated TRIM32 captures membrane-associated SseK3 for ubiquitination and subsequent proteasome degradation, which restricts SseK3-SNARE-mediated SIF biogenesis and inhibits the intracellular replication of *Salmonella* (**Figure 8**).

## Materials and Methods

### Bacterial strains, cell culture and infection

*S.* Typhimurium strains (wild-type SL1344 and its mutant derivatives) used in this study were listed in *SI Appendix,* Table S1. pET28a vector-based complementation plasmids were introduced into *S*. Typhimurium by electroporation (2.5 kV, 200 Ω, 25 μF and 5 ms). Bacteria were cultured in LB Broth at 37 °C on a shaker (220 rpm/min). When necessary, cultures were supplemented with antibiotics at the following final concentrations: streptomycin, 100 μg mL^-1^; ampicillin, 100 μg mL^-1^; kanamycin, 50 μg mL^-1^.

The procedure for bacterial infection of mammalian cells was performed as previously described. Briefly, wild-type and mutant *Salmonella* were cultured overnight (approximately 16 h) at 37 °C on a shaker (220 rpm/min) and then were subcultured at 1:33 dilution in LB without antibiotics for 3 h. Bacteria were diluted in serum-free and antibiotics-free DMEM, and added to cells at a multiplicity of infection (MOI) of 100 for 30 min at 37 °C. 24-well plates were centrifuged at 700*g* for 5 min at room temperature to promote and synchronize infection. Extracellular bacteria were removed by extensive washing with PBS, and culture media was replaced with medium containing 100 μg mL^-1^ gentamicin. Cells were incubated at 37 °C, 5% CO2 for a further 1.5 h, and the culture medium was replaced with a medium containing 20 μg mL^-1^ gentamicin. Infected cells were incubated to the indicated time at 37 °C in a 5% CO2 incubator. Samples were processed further for immunoprecipitation or immunofluorescence.

To measure the *S*. Typhimurium replication fold, RAW264.7 cells were infected with indicated *Salmonella* strains at an MOI of 10. Infection was facilitated by centrifugation at 700 g for 5 min at room temperature. After 30 min incubation at 37 °C, cells were washed three times with PBS to remove extracellular bacteria and incubated with fresh DMEM containing 100 μg mL^-1^ gentamycin. At 2 h post-infection, the gentamicin concentration was reduced to 20 μg mL^-1^. At 2 h and 24 h post-infection, cells were lysed in cold PBS containing 1% Triton X-100, and colony-forming units were determined by serial-dilution plating on agar plates containing 100 μg mL^-1^ streptomycin and 50 μg mL^-1^ kanamycin. The replication fold was determined by dividing the number of intracellular bacteria at 24 h by the number at 2 h.

### Plasmids, antibodies and reagents

Plasmids used in this study were listed in *SI Appendix,* Table S2. Genes coding for SseK1, SseK2, and SseK3 DNA were amplified from genomic DNA of *S*. Typhimurium strain SL1344 and were inserted into pCS2-EGFP, pCS2-RFP, pCS2-Flag, and pCS2-HA, respectively for transient expression in mammalian cells. For complementation in *S*. Typhimurium Δ*sseK1/2/3* strain, DNA fragment containing genes encoding SseK1, SseK2, and SseK3, each together with their upstream promoter regions, was amplified from *S*. Typhimurium SL1344 genomic DNA and inserted into pET28A. cDNAs for SNAP23, SNAP25, VAMP8, Syntaxin7, Syntaxin8, Vti1b, Sec22b, Snapin, Rab1 and TRM32 were amplified from a cDNA library of HeLa cells. For mammalian expression, cDNAs were cloned into pCS2-EGFP and pCS2-Flag vectors. Truncation, deletion, and point-mutation mutants were constructed by the standard PCR cloning strategy. All plasmids were verified by sequencing analysis. Antibodies and reagents were listed in *SI Appendix,* Table S3.

### Cell culture, transfection and stable cell-line construction

293T, HeLa cells were obtained from the American Type Culture Collection (ATCC) and were maintained in DMEM (HyClone) supplemented with 10% FBS (Gibco), 2 mM L-glutamine, 100U mL^-1^ penicillin, and 100 μg mL^-1^ streptomycin. Cells were cultivated in a humidified atmosphere containing 5% CO2 at 37 °C.

Transient transfection was performed using Vigofect (Vigorus) or Jetprime (Polyplus) reagents following the manufacturers’ instructions. For siRNA knockdown, 200 pmol of siRNAs were transfected into 2×10^6^ RAW264.7 cells. Sense sequences for the siRNAs used are as follows: *trim32* 1# 5’-CCATCTGCATGGAGTCCTTTT -3’, *trim32* 2#: 5’-CCAAGTGTTCAACCGCAAATT-3’, *trim32* 3#: 5’-GCTATCAT CTGAGAAGATATT-3’, and negative control (NC): 5’-TTCTCCGAACGTGTCAC GT-3’.

To generate the cell line that stably expresses EGFP-VAMP8, pcDNA4-EGFP-VAMP8 was transfected into 293T cells. Cell emitting green fluorescence obtained by fluorescence-activated cell sorting were cultured in DMEM medium supplemented with 10% FBS, 1% v/v penicillin/streptomycin, and 50 μg mL^-1^ zeocin. Knockout cell lines were generated by the CRISPR-Cas9 method as previously described (ref). Briefly, the pHKO plasmid containing the guide RNA targeting *Trim32* was co-transfected with packaging plasmid psPAX2 and envelope plasmid pMD2.G into Cas9-expressed 293T cell. The transfection cocktail was removed after 6 h and replaced by fresh medium. After 72 h, viral containing supernatant was collected and filtered with 0.45 μm membrane, and stored at 4 °C before transduction. 293T cells were transduced with the lentiviral particles. Three days later, GFP-positive cells were sorted into single clones in 96-well plates by flow cytometry and knockout lines were identified by PCR and by Western blot with antibodies specific for TRIM32. The sequence for the guide RNA used for *trim32* knockout is 5′-CCAGTTTGTAGTAACCGATG-3′.

### Immunoprecipitation

For immunoprecipitation, 293T cells at a confluency of 60-70% in 6-well plates were transfected with a total of 5 μg plasmids that code for the protein of interest. Twenty-four hours after transfection, cells were washed once with PBS and lysed in buffer A containing 25 mM Tris-HCl, pH 7.5, 150 mM NaCl, 10% glycerol, and 1% Triton X-100, supplemented with a protease inhibitor mixture (Roche Molecular Biochemicals). Pre-cleared lysates were subjected to anti-Flag M2 or anti-GFP immunoprecipitation following the manufacturer’s instructions. The beads were washed four times with lysis buffer, and the immunoprecipitates were eluted by SDS sample buffer followed by standard immunoblotting analysis. All the immunoprecipitation assays were performed more than three times, and representative results were shown. For enrichment of the arginine-GlcNAcylated proteins from lysates of transfected cells, samples were washed three times in ice-cold PBS and lysed in buffer A containing 25 mM Tris-HCl, pH 7.5, 150 mM NaCl, 10% glycerol, and 1% Triton X-100, supplemented with a protease inhibitor mixture (Roche Molecular Biochemicals). Pre-cleared lysates were subjected to immunoprecipitation with the anti-Arg-GlcNAc antibodies(Pan et al., 2014). The beads were washed four times with the lysis buffer, and the immunoprecipitates were dissolved by SDS sample buffer.

### Immunofluorescence labeling and confocal microscopy

At the indicated time point post-transfection or bacterial infection, cells were fixed for 10 min with 4% PFA in PBS and permeabilized for 15 min with 0.2% Triton X-100 in PBS. After blockade of nonspecific binding by incubation of cells for 30 min with 2% bovine serum albumin (BSA) in PBS, coverslips were incubated with the appropriate primary antibodies and subsequently with fluorescein-labeled secondary antibodies (ThermoFisher). Confocal fluorescence images were acquired at the confocal microscope (Spinning Disc, Leica). All image data shown are representative of at least three randomly selected fields.

### Expression and purification of recombinant proteins

Protein expression was induced in *E. coli* BL21(DE3) strain (Novagen) harboring the appropriate plasmid that directs the protein of interest at 22°C for 15 h with 0.4 mM isopropyl-β-D-thiogalactopyranoside (IPTG) after the cultures have reached an OD_600_ of 0.8–1.0. Affinity purification of GST-SseK3 was performed using glutathione sepharose (GE Healthcare), and purification of 6×His-SUMO-hUBE1, 6×His-TRIM32, 6×His-TRIM32 (ΔRING), and 6×His-Flag-Ub was conducted using Ni-NTA agarose (Qiagen), following the manufacturer’s instructions. Proteins were concentrated in a buffer containing 20 mM HEPES pH 7.5, 150mM NaCl, and 5% glycerol. The protein concentration was determined by the Bradford method (ref).

### *In vitro* ubiquitination assays

To assay ubiquitination of SseK3 *in vitro,* Ubiquitin (5 μg), E1 (200 ng), UBCH5a (300 ng), His-TRIM32 (0.8 μg) and GST-SseK3 (2 μg) were incubated with 2 mM ATP at 37°C for 2 h in ubiquitin assay buffer (20 mM Tris-HCl pH 7.5, 5 mM MgCl_2_, 2 mM DTT). Reactions were stopped by adding 30 mM EDTA and 15 mM DTT. After GST pull-down, the sample was washed with 1 M urea for 60 min to exclude potential binding of unanchored polyubiquitin, then the sample was placed in an SDS-loading buffer and boiled at 95°C for 5 min. Samples were subsequently analyzed by SDS-PAGE followed by Western blotting.

### Mass spectrometric analyses

For identification of the GlcNAcylated arginine and arginine-containing peptides, purified SNAP25 protein was subjected to digestion with GluC and LysC, and the resulting peptides were separated on an EASY-nLC 1200 system (Thermo Fisher Scientific). The nano liquid chromatography gradient was as follows: 0-8% B in 3 min, 8-28% B in 42 min, 28-38% B in 5 min, 38-100% B in 10 min (solvent A: 0.1% Formic acid in water, solvent B: 80% CH3CN in 0.1% formic acid). Peptides eluted from the capillary column were applied directly onto a Q Exactive Plus mass spectrometer by electrospray (Thermo Fisher Scientific) for mass spectrometry (MS) and MS/MS analyses. Searches were performed against the amino acid sequence of SNAP25 and were performed with cleavage specificity allowing four mis-cleavage events. Searches were performed with the variable modifications of oxidation of methionine, *N*-Acetyl-hexosamine addition to arginine (Arg-GlcNAc), and acetylation of protein N termini. For identification of the SNAP25-binding protein, immunoprecipitates were separated using SDS-PAGE, fixed, and visualized after silver staining as recommended by the manufacturer. An entire lane of bands was excised and subjected to in-gel trypsin digestion and MS/MS detections as described above. Identification of proteins was carried out using the Proteome Discoverer 2.2 program. Searches were performed against the Human proteomes depending on the samples with carbamidomethylation of cysteine set as a fixed modification. The precursor mass tolerance was set to 10 parts-per-million (ppm) and a fragment mass tolerance of 0.02 Da. A maximum false discovery rate (FDR) of 1.0% was set for protein and peptide identifications.

### Quantification and statistical analysis

All results are presented as mean ± standard deviation containing a specified number of replicates. Data were analyzed using a Student’s *t*-test to compare two experimental groups. The comparison of multiple groups was conducted by using the one-way analysis of variance (ANOVA). A difference is considered significant as the following: \**P* <0.05, \*\**P* < 0.01.

## Compliance and ethics

All authors declare no competing interests. This study does not involve human subjects and animals.

## Author Contributions

S.L. and K.M. conceived the study. K.M. and J.Y. designed and performed the functional experiments. K.M. and J.X. conducted the mass spectrometry experiments. L.S. provided technical assistance in the analyses of the mass spectrometry data. J.L. and P.Z. provided assistance in preparing experiments materials. S.L. and K.M. analyzed the data and wrote the manuscript. All authors discussed the results and commented on the manuscript.

## Acknowledgments

We thank members of the Li laboratory and the central laboratory of Taihe hospital for helpful discussions and technical assistance. We thank Prof. Hongbing Shu and Prof. Bo Zhong in Wuhan University (China) for providing the pRK-HA-Ub (K48 only), and pRK-HA-Ub (K63 only) plasmids. We thank Prof. Gang Cao in Huazhong Agricultural University (China) for providing the pHKO14-Cas9-Flag, and pHKO-GFP-sgRNA plasmids. This work was supported by the National Key Research and Development Programs of China 2021YFD1800404 and 2018YFA0508000, Huazhong Agricultural University Scientific & Technological Self-Innovation Foundation 2017RC003 to S.L., and Hubei Provincial Natural Science Foundation 2021CFB472 to K.M..

## Supplementary Information for

### SupplementaryTables

**Table S1.** Bacterial strains used in this study.

| Strain or plasmid | Relevant characteristics | References |
| --- | --- | --- |
| BL21(DE3) | used for protein expression/purification | Novagen |
| DH5α | used for cloning | Novagen |
| SL1344 | wild type *S.Typhimurium*, Strep<sup>r</sup> | ATCC |
| SL1344 Δ*ssaV* | *ssaV* gene deleted in SL1344, Strep<sup>r</sup> | [1] |
| SL1344 Δ*sseK1/2/3* | *sseK1*, *sseK2* and *sseK3* gene deleted in SL1344, Strep<sup>r</sup> | [1] |
| SL1344 Δ*sseK1/2/3*+pET28a | Δ*sseK1/2/3* containing pET28a, Strep<sup>r</sup>, Km<sup>r</sup> | [1] |
| SL1344 Δ*sseK1/2/3*+pSseK1 | Δ*sseK1/2/3* containing pET28a-SseK1, Strep<sup>r</sup>, Km<sup>r</sup> | [1] |
| SL1344 Δ*sseK1/2/3*+pSseK2 | Δ*sseK1/2/3* containing pET28a-SseK2, Strep<sup>r</sup>, Km<sup>r</sup> | [1] |
| SL1344 Δ*sseK1/2/3*+pSseK3 | Δ*sseK1/2/3* containing pET28a-SseK3, Strep<sup>r</sup>, Km<sup>r</sup> | [1] |
| SL1344 Δ*sseK1/2/3*+pSseK3 DxD | Δ*sseK1/2/3* containing pET28a-SseK3 DxD, Strep<sup>r</sup>, Km<sup>r</sup> | [1] |

**Table S2.** Plasmids used in this study.

| Plasmids | Relevant characteristics | References |
| --- | --- | --- |
| pCS2-GFP-SseK3 | pCS2-GFP carrying SseK3 coding region, Amp<sup>r</sup> | [1] |
| pCS2-GFP-SseK3 Dx D | pCS2-GFP carrying SseK3 D226A/D228A mutation, Amp<sup>r</sup> | [1] |
| pCS2-RFP-SseK3 | pCS2-RFP carrying SseK3 coding region, Amp<sup>r</sup> | [1] |
| pCS2-Flag-SNAP23 | pCS2-Flag carrying SNAP23 coding region, Amp<sup>r</sup> | This study |
| pCS2-Flag-SNAP25 | pCS2-Flag carrying SNAP25 coding region, Amp<sup>r</sup> | This study |
| pCS2-GFP-SNAP25 | pCS2-GFP carrying SNAP25 coding region, Amp<sup>r</sup> | This study |
| pCS2-Flag-SNAP25 NTD | pCS2-Flag carrying SNAP25 (1-90 aa) region, Amp<sup>r</sup> | This study |
| pCS2-HA-SNAP23 | pCS2-HA carrying SNAP23 coding region, Amp<sup>r</sup> | This study |
| pCS2-HA-SNAP25 | pCS2-HA carrying SNAP25 coding region, Amp<sup>r</sup> | This study |
| pCS2-Flag-VAMP8 | pCS2-Flag carrying VAMP8 coding region, Amp<sup>r</sup> | This study |
| pCS2-HA-VAMP8 | pCS2-HA carrying VAMP8 coding region, Amp<sup>r</sup> | This study |
| pcDNA4-GFP-VAMP8 | pcDNA4 carrying GFP-VAMP8 coding region, Amp<sup>r</sup> | This study |
| pCS2-Flag-Vti1b | pCS2-Flag carrying Vti1b coding region, Amp<sup>r</sup> | This study |
| pCS2-Flag-Sec22b | pCS2-Flag carrying Sec22b coding region, Amp<sup>r</sup> | This study |
| pCS2-Flag-Snapin | pCS2-Flag carrying Snapin coding region, Amp<sup>r</sup> | This study |
| pCS2-Flag-Syntaxin 7 | pCS2-Flag carrying Syntaxin7 coding region, Amp<sup>r</sup> | This study |
| pCS2-Flag- Syntaxin 8 | pCS2-Flag carrying Syntaxin8 coding region, Amp<sup>r</sup> | This study |
| pCS2-Flag- Rab1 | pCS2-Flag carrying Rab1 coding region, Amp<sup>r</sup> | [1] |
| pCS2-Flag-TRIM32 WT | pCS2-Flag carrying TRIM32 (1-653aa) region, Amp<sup>r</sup> | This study |
| pCS2-Flag-TRIM32 RING | pCS2-Flag carrying TRIM32 (1-96 aa) region, Amp<sup>r</sup> | This study |
| pCS2-Flag-TRIM32 RING+B-box | pCS2-Flag carrying TRIM32 (1-135 aa) region, Amp<sup>r</sup> | This study |
| pCS2-Flag-TRIM32 NHL | pCS2-Flag carrying TRIM32 (255-653 aa) region, Amp<sup>r</sup> | This study |
| pCS2-Flag-TRIM32 ΔRING | pCS2-Flag carrying TRIM32 (97-653 aa) region, Amp<sup>r</sup> | This study |
| pCS2-Flag-TRIM32 Δ(RING+B-box) | pCS2-Flag carrying TRIM32 (136-653 aa) region, Amp<sup>r</sup> | This study |
| pCS2-Flag-TRIM32 ΔNHL | pCS2-Flag carrying TRIM32 (1-254 aa) region, Amp<sup>r</sup> | This study |
| pCS2-Flag-TRIM32 C39S | pCS2-Flag carrying TRIM32 C39S mutation, Amp<sup>r</sup> | This study |
| pCS2-Flag-TRIM32 R394H | pCS2-Flag carrying TRIM32 R394H mutation, Amp<sup>r</sup> | This study |
| pCS2-Flag-TRIM32 D487N | pCS2-Flag carrying TRIM32 D487N mutation, Amp<sup>r</sup> | This study |
| pHKO14-Cas9 | SpCas9 expression plasmid, blasticidin resistance | Gang Cao's lab |
| pHKO-GFP-sgRNA | sgRNA cloning backbone for expression of sgRNA in mammalian cells, puromycin resistance | Gang Cao's lab |
| pRK5-HA-Ub WT | pRK5-HA carrying wild type ubiquitin | Addgene |
| pRK5-HA-Ub K27R | pRK5-HA carrying ubiquitin K27R mutation | This study |
| pRK5-HA-Ub K29R | pRK5-HA carrying ubiquitin K29R mutation | This study |
| pRK5-HA-Ub K33R | pRK5-HA carrying ubiquitin K33R mutation | This study |
| pRK5-HA-Ub K48R | pRK5-HA carrying ubiquitin K48R mutation | This study |
| pRK5-HA-Ub K63R | pRK5-HA carrying ubiquitin K63R mutation | This study |
| pRK5-HA-Ub K48 only | all Lys residues are mutated to Arg except K48 | [2] |
| pRK5-HA-Ub K63R only | all Lys residues are mutated to Arg except K63 | [2] |

**Table S3.** Antibodies and reagents used in this study.

| REAGENT or RESOURCE | SOURCE | IDENTIFIER |
| --- | --- | --- |
| **Antibodies** |  |  |
| Mouse monoclonal anti-tubulinA | Sigma-Aldrich | Cat#T5168 |
| Rabbit polyclonal anti-Arg-GlcNAc | Abcam | Cat#ab195033 |
| Mouse monoclonal anti-Flag | Sigma-Aldrich | Cat#F7425 |
| Mouse monoclonal anti-EGFP | Santa Cruz Biotechnology | Cat#sc8334 |
| Mouse monoclonal anti-HA | Biolegend | Cat#901501 |
| Rabbit polyclonal anti-Salmonella | Abcam | Cat#ab35156 |
| Mouse monoclonal anti-Ubiquitin P4D1 | Santa Cruz Biotechnology | Cat#sc8017 |
| Mouse monoclonal anti-ATP1B1 | Santa Cruz Biotechnology | Cat#sc21713 |
| Mouse monoclonal anti-Tom20 | Santa Cruz Biotechnology | Cat#sc17764 |
| Mouse monoclonal anti-GM130 | BD Biosciences | Cat#5239872 |
| Goat Anti-Rabbit IgG H&L (Alexa Fluor® 488) | Thermo Fisher Scientific | A32731 |
| Goat Anti-Mouse IgG H&L (Alexa Fluor® 488) | Thermo Fisher Scientific | A32723 |
| Goat Anti-Rabbit IgG H&L (Alexa Fluor® 564) | Thermo Fisher Scientific | A-11035 |
| Goat Anti-Mouse IgG H&L (Alexa Fluor® 564) | Thermo Fisher Scientific | A-11030 |
| Goat Anti-Rabbit IgG H&L (Alexa Fluor® 647) | Thermo Fisher Scientific | A32733 |
| Goat Anti-Mouse IgG H&L (Alexa Fluor® 647) | Thermo Fisher Scientific | A32728 |
| **Chemicals, peptides, and recombinant proteins** |  |  |
| DAPI | Sigma-Aldrich | Cat#D9542 |
| MG-132 | Sigma-Aldrich | Cat#C2211 |
| Chloramphenicol | Sigma-Aldrich | Cat#R4408 |
| Isopropyl β-D-Thiogalactoside (IPTG) | Sigma-Aldrich | Cat#I6758 |
| Cycloheximide | Sigma-Aldrich | Cat#C7698 |
| Ampicillin | Sigma-Aldrich | Cat#A9393 |
| Gentamicin | Sigma-Aldrich | Cat#G1397 |
| Kanamycin | Sigma-Aldrich | Cat#K1377 |
| Dulbecco's Modified Eagle's Medium, High Glucose | Gibco | Cat#C11885500BT |
| PBS pH 7.4 basic (1×) | Gibco | Cat#C10010500BT |
| Fetal Bovine Serum | Biological Industries | Cat#04-001-1ACS |
| 0.05% Trypsin-EDTA | Thermo Fisher Scientific | Cat#25300054 |
| Complete Protease Inhibitor | Roche | Cat#11836170001 |
| DAPI | Thermo Fisher Scientific | Cat#D1306 |
| jetPRIME® | Polyplus transfection | Cat#114-15 |
| VigoFect | Vigorous biotechnology | Cat#T001 |
| Trypsin Protease, MS Grade | Thermo Fisher Scientific | Cat#90057 |
| Lys-C Endoproteinase, MS Grade | Thermo Fisher Scientific | Cat#90051 |
| Glu-C Endoproteinase | Thermo Fisher Scientific | Cat#90054 |
| EZview™ Red® ANTI-FLAG M2 Affinity Gel | Sigma-Aldrich | Cat#F2426 |
| Anti-GFP Affinity Beads 4FF | SMART LIFESCIENCES | Cat#SA070001 |

---

**Figure S1.**
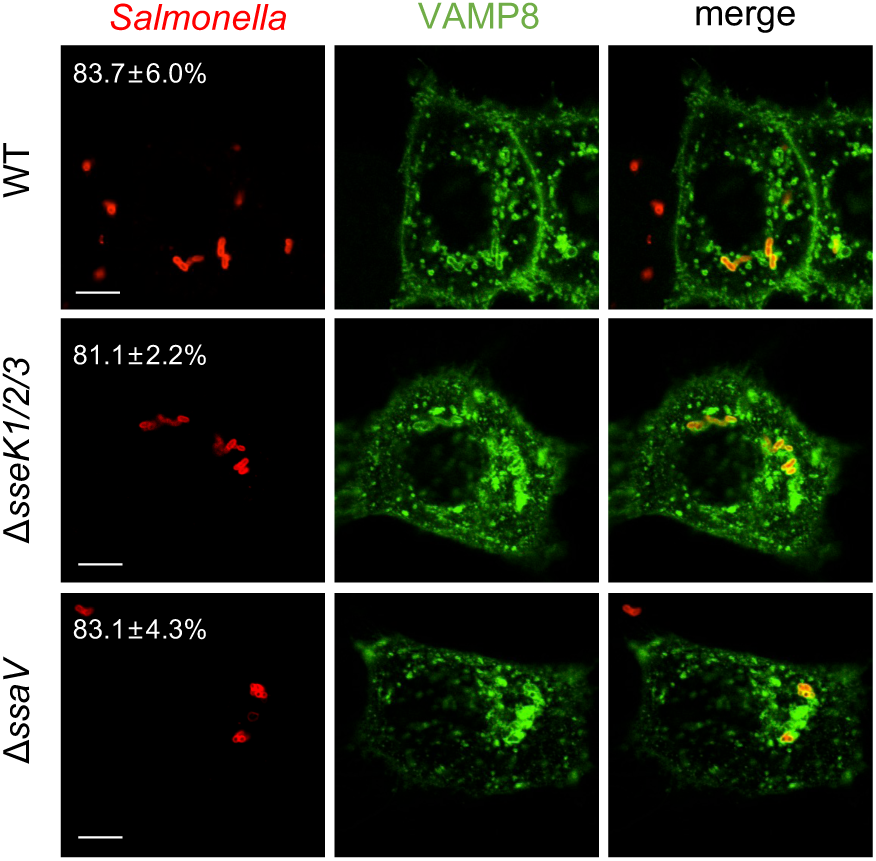
The Effects of SseKs on the frequency of VAMP-8 coated *Salmonella*. EGFP-VAMP8 stably expressed HeLa cells were infected with the indicated *Salmonella* strains for 2 hr. Shown are fluorescence detection of VAMP8 (green) and *Salmonella* (red). Statistics of cells showing the frequency of VAMP-8 coated *Salmonella* are listed in the upper left corner. At least 100 cells were counted for each experiment, and the statistical data shown are from three independent determinations. Scale bar, 10 μm.

**Figure S2.**
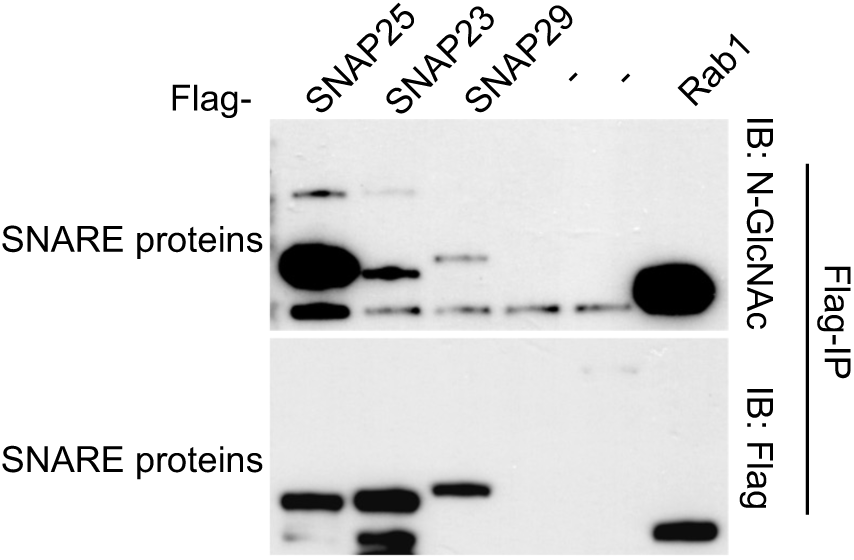
Modification of SNAP proteins by SseK3 during *S*. Typhimurium infection. 293T cells were transfected with a plasmid expressing the Flag-SNAP23, Flag-SNAP25, Flag-SNAP29 or Flag-Rab1 individually, and then infected with *S.* Typhimurium Δ*sseK1/2/3* complemented with pET28a-SseK3. After infection, cells were lysed, and proteins were immunoprecipitated with anti-Flag beads, followed by standard immunoblotting analysis with the indicated antibodies.

**Figure S3.**
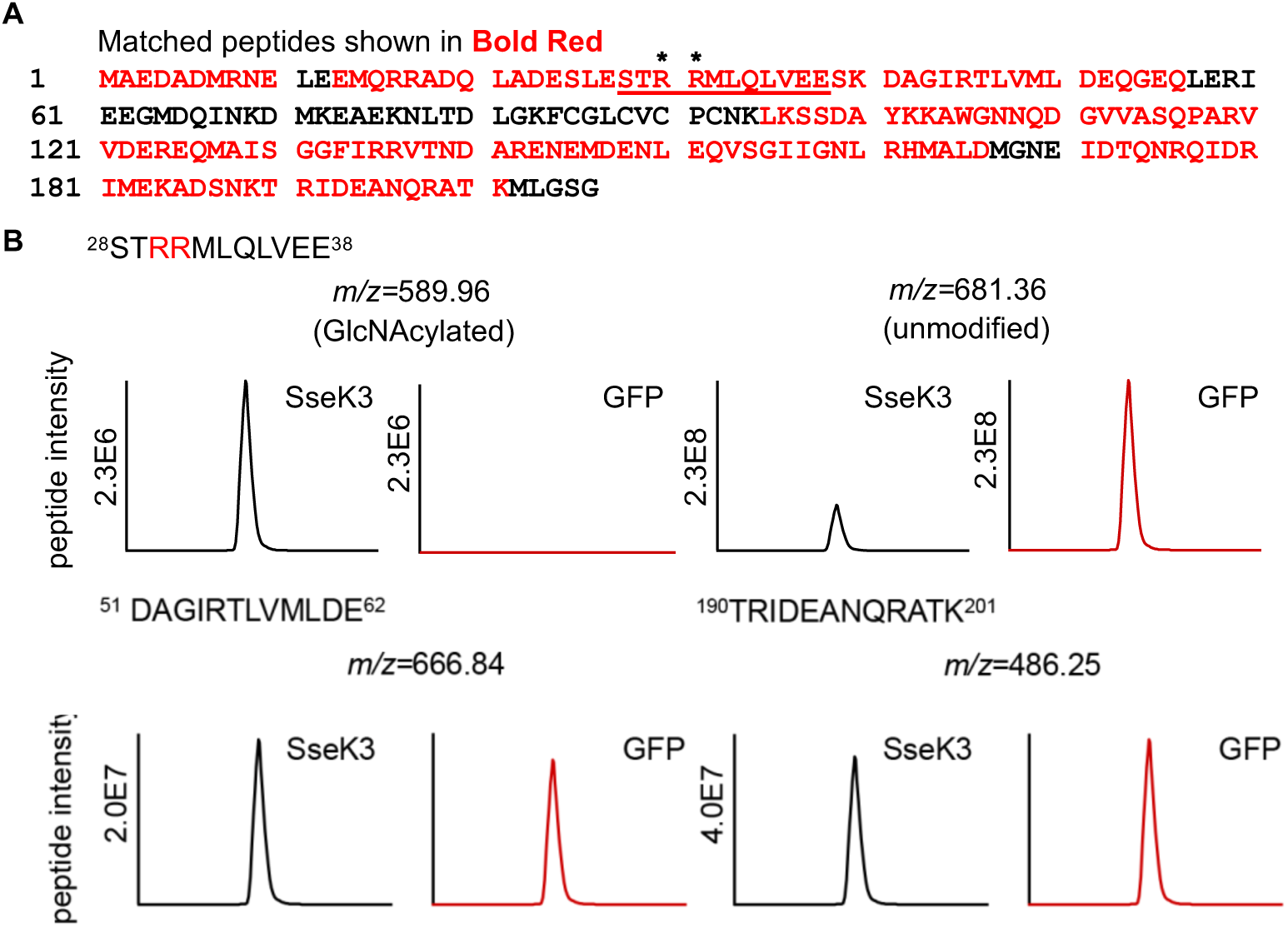
MS detection of SNAP25 peptides intensity. (A) Flag-SNAP25 was isolated from 293T cells co-transfected with either wild-type GFP-SseK3 or the empty plasmid GFP. Immunoprecipitated SNAP25 was then digested with GluC and LysC, and analyzed by LC-MS/MS. Detected SNAP25 sequence shown in red in LC-MS experiments. (B) The GlcNAcyalted peptide sequence is underlined, and asterisks indicate the modification. The peptide ^28^STRRMLQLVEE^38^ covalently modified with GlcNAc is shown, and the detected arginines are labeled in red. Two control peptides are also displayed. Extracted ion chromatograms of the doubly protonated peptide are shown with peak intensities indicating the relative amounts of either the modified or unmodified peptides. These data correspond to Fig 2.

